# Intrinsic strain-specific behaviour predicts emergent collective aggregation in heterogeneous *C. elegans* groups

**DOI:** 10.1101/2025.10.06.680706

**Authors:** Narcís Font-Massot, Jacob D. Davidson, Siyu Serena Ding

**Affiliations:** Centre for the Advanced Study of Collective Behaviour, 78464 Konstanz, Germany; Max Planck Institute of Animal behaviour, 78464 Konstanz, Germany; International Max Planck Research School for Quantitative Behaviour, Ecology and Evolution, 78464 Konstanz, Germany; Department of Mathematics and Computer Science, Freie Universität Berlin, 14195 Berlin, Germany

**Keywords:** Collective behaviour, Heterogeneity, *C. elegans*, Aggregation

## Abstract

Collective animal behaviour research to date typically specifies members of the group as identical individuals, even though within group heterogeneity is commonplace. We exploit the tractable *C. elegans* study system to explicitly define and manipulate heterogeneity to investigate how individuals with different behavioural phenotypes interact and aggregate in heterogeneous group settings. Using controlled mixing experiments between pairs of strains that have defined aggregation tendencies, we apply a quantitative behavioural analysis framework and show that individuals maintain their intrinsic movement patterns and interaction rules regardless of group composition. Notably, neither behavioural differences nor distant genetic relatedness between strains lead to a modulation of individual behaviour; instead, distinct strains behave and coexist without influencing each other’s intrinsic behavioural tendencies. Using a simulation model, we further show that aggregation in mixed *C. elegans* groups can be accurately predicted from strain-specific individual-level parameters measured in homogeneous settings. Our integrated approach provides a generalised framework for understanding collective behaviour in diverse heterogeneous systems, which may offer insights into population-level consequences of phenotypic variation and broader ecological processes.

## 1. Introduction

Collective behaviour emerges from interactions among individuals, yet research to date has predominantly treated these individuals as identical [1, 2]. This oversimplification does not usually reflect biological reality, neglecting the inherent individual differences often present in such systems. Recent work has shown how heterogeneity – driven by genetic, physiological and informational differences between individuals – plays a crucial role in the emergence and functionality of collectives [3, 4, 5, 6, 7] and influences broader ecological patterns and processes [8, 9]. However, there is a notable lack of work that explicitly defines and manipulates heterogeneity inside groups to quantitatively probe how individual behaviour and inter-individual interactions shape collective behaviour in these settings.

To this end, we introduce the nematode *Caenorhabditis elegans* as an experimental system for investigating the effect of heterogeneity on collective behaviour. This model organism species has been extensively studied from genetic, neuronal, and behavioural perspectives. We can leverage this vast knowledge base to precisely control the behavioural characteristics and relatedness between individuals to answer our questions about the mechanisms of heterogeneous collective behaviour. At the same time, this line of enquiry also has potential implications for understanding the ecology of this species. Previous work has shown that clonal groups of *C. elegans* show diverse aggregation behaviours ranging from solitary to strongly aggregating on a food patch [10, 11, 12]. While rapidly proliferating *C. elegans* populations on resource patches are likely clonal in nature due to the self-fertilizing mode of reproduction, this species also engages in long-range phoretic dispersal leading to local genetic diversity [13, 14, 15]. Despite reports that multiple strains can co-occur in nature and exploit the same resource patch during population growth [16, 17], it is not known how heterogeneous groups consisting of different *C. elegans* strains interact together on a food patch.

In this work, we establish an experimental, analytical and modelling framework to study heterogeneous *C. elegans* groups and apply this framework to investigate the motility and interaction mechanisms underlying the collective behaviour of mixed groups. We perform two sets of experiments that each mix together two different *C. elegans* strains, where the aggregation behaviour phenotype for each strain in homogeneous group settings is known. In the first set of experiments (MIX-1), we combine a solitary strain with a genetically-related aggregating strain to ask if and how individuals with different aggregation tendencies may influence each other’s behaviour. In the second set (MIX-2), we mix two distantly related strains with similar aggregation tendencies to ask whether unrelated individuals form hybrid aggregates, and which motility and interaction mechanisms underlie this process. For both MIX-1 and MIX-2, we use a battery of metrics to quantify motion and aggregation, while explicitly considering neighbour identity and the response to both same-type and other-type individuals. To interpret these results, we develop a simple simulation model of worm collective behaviour. We examine whether the model can reproduce experimental results for homogeneous cases and then use simulations to evaluate its predictive ability for mixed groups.

## 2. Results

### 2.1 Phenotypically different *C. elegans* strains do not influence each other’s behaviour in heterogeneous groups

Previous work in collective behaviour research has shown that when two different types of individuals interact, those that are normally solitary can start to behave more socially [18, 19]; furthermore, theoretical studies have shown that differences in individual behaviour can influence group cohesion [20, 21]. In the first set of experiments (MIX-1), we tested whether two *C. elegans* strains with different aggregation tendencies influence each other when mixed in a heterogeneous group. Specifically, we used the solitary laboratory reference strain N2 and the aggregating strain *npr-1* [10], and asked whether mixing causes N2 worms to disrupt the cohesion of *npr-1* aggregates, or instead to join them.

To answer these questions, we compared homogeneous groups consisting of 40 individuals of a single strain with heterogeneous groups composed of 20 individuals from each of the two strains. The individuals freely behaved on a 1 cm (10 times the individual’s body length) diameter round *Escherichia coli* OP50 food patch, and were continuously recorded for 45 minutes at 10 fps. We used fluorescence pharyngeal muscle markers to identify individuals belonging to each strain: N2 with a red marker and *npr-1* with a green marker (Fig. 1a) (see Table 1 for a list of strains and genotypes), and performed automated tracking of fluorescence signals [22] to obtain the motion trajectories and strain identity during each trial for precise behaviour quantification.

**Table 1:** List of *C. elegans* strain names, description, genotype and source used in this work.

| Strain name | Description | Genotype | Source |
| --- | --- | --- | --- |
| N2 | Unlabelled N2 | Lab reference strain | *Caenorhabditis* Genetics Center (CGC) |
| ATU4301 | Red labelled N2, referred to in this study as "N2" | *aceIs1 [myo-3p::mitochondrial LAR-GECO + myo-2p::RFP]* in N2 background | Higashitani Lab |
| DA609 | Unlabelled *npr-1* | *npr-1(ad609)X* in N2 background | CGC |
| OMG2 | Green labelled *npr-1*, referred to in this study as "*npr-1*" | *mIs12[myo-2p::GFP+pes-10p::GFP+F22B7.9p::GFP]II;npr-1(ad609)X* in N2 background | Brown Lab |
| CB4856 | Unlabelled CB4856 | Hawaiian wild isolate | CGC |
| SSD04 | Red labelled CB4856, referred in this study as "CB4856" | *aceIs1[myo-3p::mitochondrial LAR-GECO + myo-2p::RFP]* in CB4856 background, introgressed 10x | This study |

**Figure 1:**
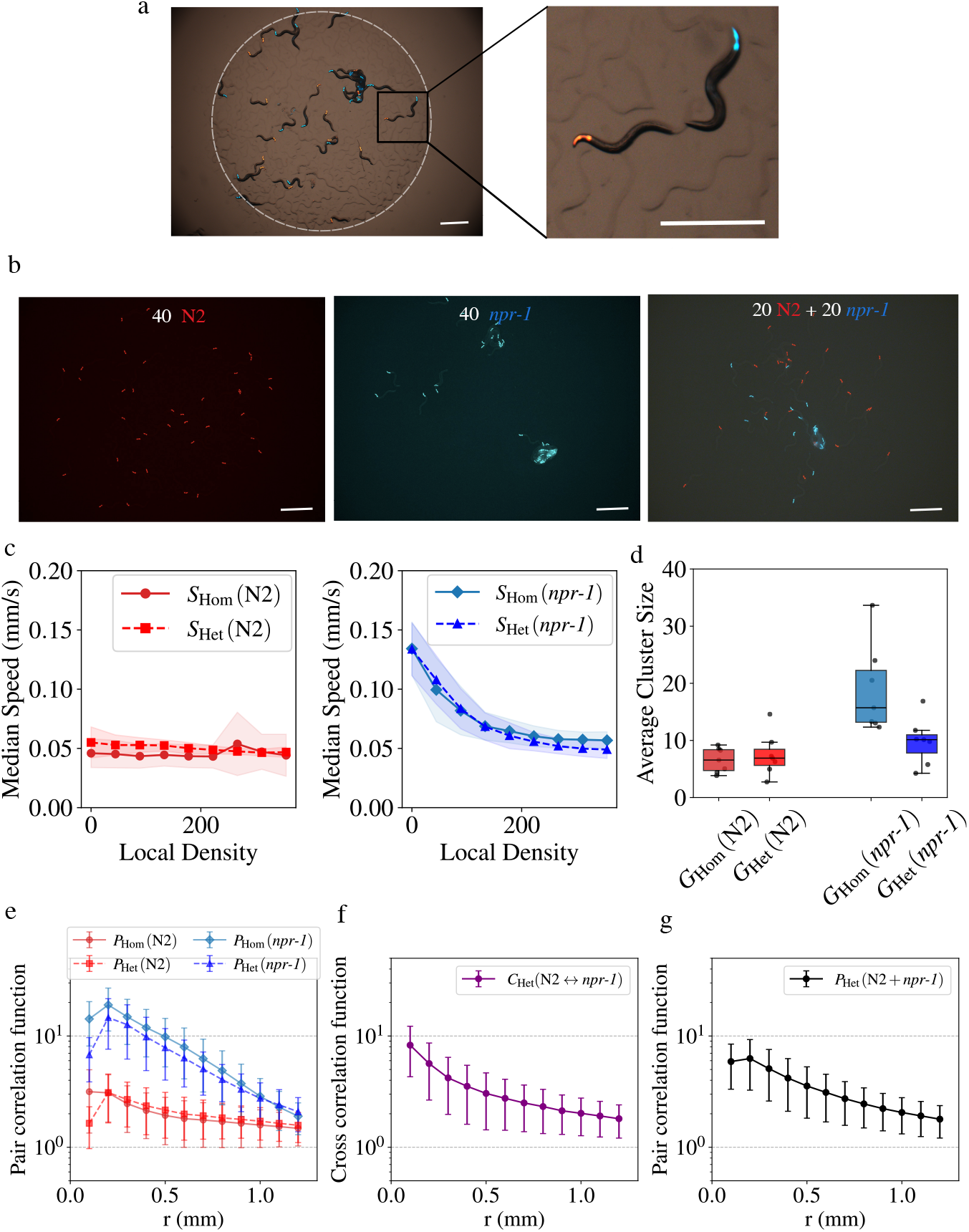
Behaviour quantification for MIX-1 experiments. **(a)** (Left) Snapshot of an experiment with two different worm strains interacting in a 1 cm diameter OP50 bacteria food arena (dashed white contour). (Right) Zoom-in of the experiment showing the two different *C. elegans* strains labeled with red and green fluorescence markers for strain identification and tracking. Scale bars = 1 mm. **(b)** Representative snapshots of homogeneous experiments with 40 N2 individuals (left) or 40 *npr-1* individuals (middle), and heterogeneous experiments with 20 *npr-1* and 20 N2 individuals. Scale bars = 1 mm. See Supplementary Videos 1, 2, and 4 for additional representative examples of the behaviours. **(c)** Density dependent median speed (*S*) distributions for each strain in homogeneous (*S*_Hom_; solid line) and heterogeneous (*S*_Het_; dashed line) trials. The line shading represents the standard deviation of the data around the mean. **(d)** Average group size (*G*) for N2 and *npr-1* strains in homogeneous (*G*_Hom_) and heterogeneous (*G*_Het_) trials. The boxplots show the median (central line), interquartile range (box), and non-outlier data range (whiskers), with points beyond the whiskers representing outliers. **(e)** Pair correlation function (*P*) for the homogeneous (*P*_Hom_; solid line) and heterogeneous trials (*P*_Het_; dashed line) for the N2 and *npr-1* strains. **(f)** Cross correlation function (*C*_Het_) between the two strain types. **(g)** Pair correlation function (*P*_Het_) considering all individuals in heterogeneous groups. The error bars represent the the standard deviation of the data around the mean. The sample size is n = 7 for each experimental condition.

Qualitative observations of the homogeneous trials suggested consistency with previous results [10]: N2 individuals are mostly solitary and scattered across the food patch, whereas *npr-1* worms form aggregates with individuals tightly packed together with strong physical overlap (Fig 1b, left and middle, Supp. Vid. 1-2). In heterogeneous groups, *npr-1* worms still form aggregates despite a lower strain density (now only 20 *npr-1* worms in the arena instead of 40), but the aggregates appear smaller, occasionally with N2 worms present within *npr-1* aggregates (Fig. 1b, right, Supp. Vid. 4).

Next, we performed comprehensive quantitative analyses to further describe these observations and compare the individual and collective behaviours across homogeneous and heterogeneous social environments. This analysis proceeds in four steps: first, examining individual motility and response to the local social environment; second, analysing aggregate (or cluster) properties; third, assessing the spatial organization of individuals for each strain; and finally, exploring the spatial relationships between the two strains and analysing the collective behaviour of the heterogeneous group by inspecting the spatial organization of all individuals.

### 2.2. Motility and density-dependence for each strain

Previous work on aggregate formation in *C. elegans* has shown that individual-level behaviour, particularly density-dependent speed, reorientation and reversals, plays a key role in the formation and the stability of aggregates [23]. To test whether worms alter or maintain their motility trends in heterogeneous environments, we examined median speed (*S*) as a function of local density across strains. We defined local density as the number of neighbours within a circle area where the radius is equal to one body length (the full body length of 1 mm was used here even though we only tracked fluorescent heads in our data). In homogeneous trials when worms are at zero local density (no close neighbours), speed differs across strains: *npr-1* worms show the highest S, while N2 worms have lower speeds, revealing inherent differences in motility (Fig. 1c). As local density increases, *npr-1* worms decrease their speed, whereas N2 worms show no change. Comparing heterogeneous versus homogeneous groups, the trends for S as a function of local density show no differences (Fig. 1c). This suggests that individuals do not adjust their motility based on the strain identity of nearby worms.

To further explore whether neighbour strain identity influences motility, we compared cases where an individual has more local neighbours of the same or the other strain, and found no speed differences based on local neighbour strain identity (Supp. Fig. 1). We also analysed angular velocity (*W*) as a proxy for reorientation and reversal behaviour and obtained the same results: a decrease with the local density for *npr-1*, no local density effect for N2, and no effect for either strains in relation to the local density strain composition (Supp. Fig. 1a). These findings indicate that motility is driven solely by local density rather than by the composition of the local or overall social environment.

#### 2.1.2 Cluster properties for each strain

We next examined the structure of group aggregation behaviour by defining “clusters” consisting of at least three closely aggregating individuals (see Methods for details). With this, we quantified the average cluster size (*G*) an individual of each strain belongs to. In homogeneous groups, *G*_Hom_ reveals the expected strain-specific differences: N2 worms exhibit smaller average cluster sizes, whereas *npr-1* worms form larger clusters (Fig. 1d). In heterogeneous groups, N2 worms do not form larger clusters (c.f. *G*_Hom_ and *G*_Het_ for N2 in Fig. 1d). However, *npr-1* cluster size in heterogeneous groups decreases compared to the homogeneous case (Fig. 1d).

To describe clustering in more detail, we examined the mean fraction of individuals inside clusters (*F*). While we found similar *F* values between the *Hom* and *Het* cases for N2, there is just a slight decrease in *F* for *npr-1* in the heterogenous case compared to the homogeneous case. The changes in G for *npr-1* therefore simply reflect the effect of having a reduced number of aggregating worms in the heterogeneous groups (i.e., there are half as many *npr-1* worms in the heterogenous trials compared to homogenous *npr-1* trials), rather than a disruption of *npr-1* aggregates by N2, which would be indicated by a much lower fraction of *npr-1* worms in clusters in the heterogeneous case (*F*_Het_). This highlights the ability of *npr-1* worms to form aggregates even in heterogeneous groups with fewer aggregating worms present.

#### 2.1.3 Spatial organisation for each strain

Building on the cluster metrics, we quantified further details of spatial organization using the pair correlation function *P*, which measures the probability of finding another individual at distance *r*. Higher *P* indicates aggregation-like behaviour with more neighbours at the distance *r*, whereas lower values represent more solitary organisation. Considering only spatial relationships among same-strain individuals, we saw similar spatial patterns in homogeneous (*P*_Hom_) and heterogeneous (*P*_Het_) conditions: *npr-1* worms are found near other *npr-1* worms, while N2 worms remained more dispersed (Fig. 1e). The simpler scalar metric of mean neighbour distance shows the same trends (Supp. Fig. 1c). These metrics indicate that the internal structure of aggregates for each strain do not change with heterogeneity in their social environments.

#### 2.1.4 Spatial relationship between strains

The above analysis describes each strain separately but does not address how the strains interact spatially. We therefore examined the cross correlation *C*(*r*), which measures the probability of finding an individual of the other strain at distance *r* (with *C* = 1 indicating independent spatial distributions). In heterogeneous groups, *C*_Het_(N2 → *npr-1*) lay between the same-strain pair correlations; this means that *npr-1* worms were more likely to be surrounded by other *npr-1* than by N2, yet N2 can still be found within *npr-1* aggregates (Fig. 1f). Using the additional measure of spatial density overlap, we found low but above random values, indicating that N2 worms are indeed sometimes found in *npr-1* aggregates (Supp. Fig. 1d). Further supporting this – and consistent with the cross-correlation result – the pair correlation computed for all worms in the mixed group regardless of strain identity, also falls between the two cases for the homogeneous groups (Fig. 1g). Together, these results indicate partial spatial overlap: the strains neither fully segregate nor uniformly mix, and while mixed aggregates can form, they predominantly include *npr-1* worms.

To summarize the results for MIX-1, strain-specific motility and aggregation remain unchanged in heterogeneous groups: *npr-1* decreases speed with density whereas N2 does not, and these trends are independent of neighbour identity. Mixed groups show partial spatial overlap – N2 can occur within *npr-1* aggregates, but aggregates are predominantly formed with *npr-1* worms – and the overall spatial organization is intermediate between the homogeneous cases. The smaller *npr-1* aggregates in mixed trials arise simply from having fewer *npr-1* individuals in comparison to homogeneous *npr-1* trials, and not from disruption by N2.

### 2.2 Genetically distinct *C. elegans* strains aggregate together without behavioural modulation

Previous work in collective animal behaviour has shown that genetic differences amongst members can influence group behaviour [4, 24, 25]. In the MIX-1 experiments above, although the N2 and *npr-1* strains displayed different aggregation behaviours, they are genetically nearly identical except for a single gene difference [10]. In this section, we ask whether genetic divergence between strains constrains their ability to aggregate, or whether behavioural similarity alone is sufficient for them to form mixed aggregates in a heterogeneous social environment. To address these questions, in MIX-2 experiments we combined the aggregating laboratory strain *npr-1* with the genetically distant, aggregating wild isolate CB4856 [26]. In the absence of an explicitly known kin-recognition system in *C. elegans*, we asked whether both strains would aggregate together, and if so, which mechanisms might underlie their collective behaviour. Using the same setup as in MIX-1, we labelled CB4856 with a red fluorescence marker and *npr-1* with a green marker (Table 1) to observe their collective behaviour.

An initial qualitative inspection of the data confirms the expected aggregation patterns in homogeneous groups: both *npr-1* and the Hawaiian CB4856 strain form aggregates, although CB4856 tendency to aggregate is lower (Fig. 2a, left and middle, Supp. Vid. 2-3). In heterogeneous groups, visual inspection suggests that both strains indeed form mixed aggregates together. We also observed that the aggregates are dynamic, where worms can disperse and form new aggregates (Fig. 2a, right, Supp. Vid. 5). Following the same methods as MIX-1, we next computed a series of metrics to make a detailed comparison of the motility and spatial organization of the worms for MIX-2.

**Figure 2:**
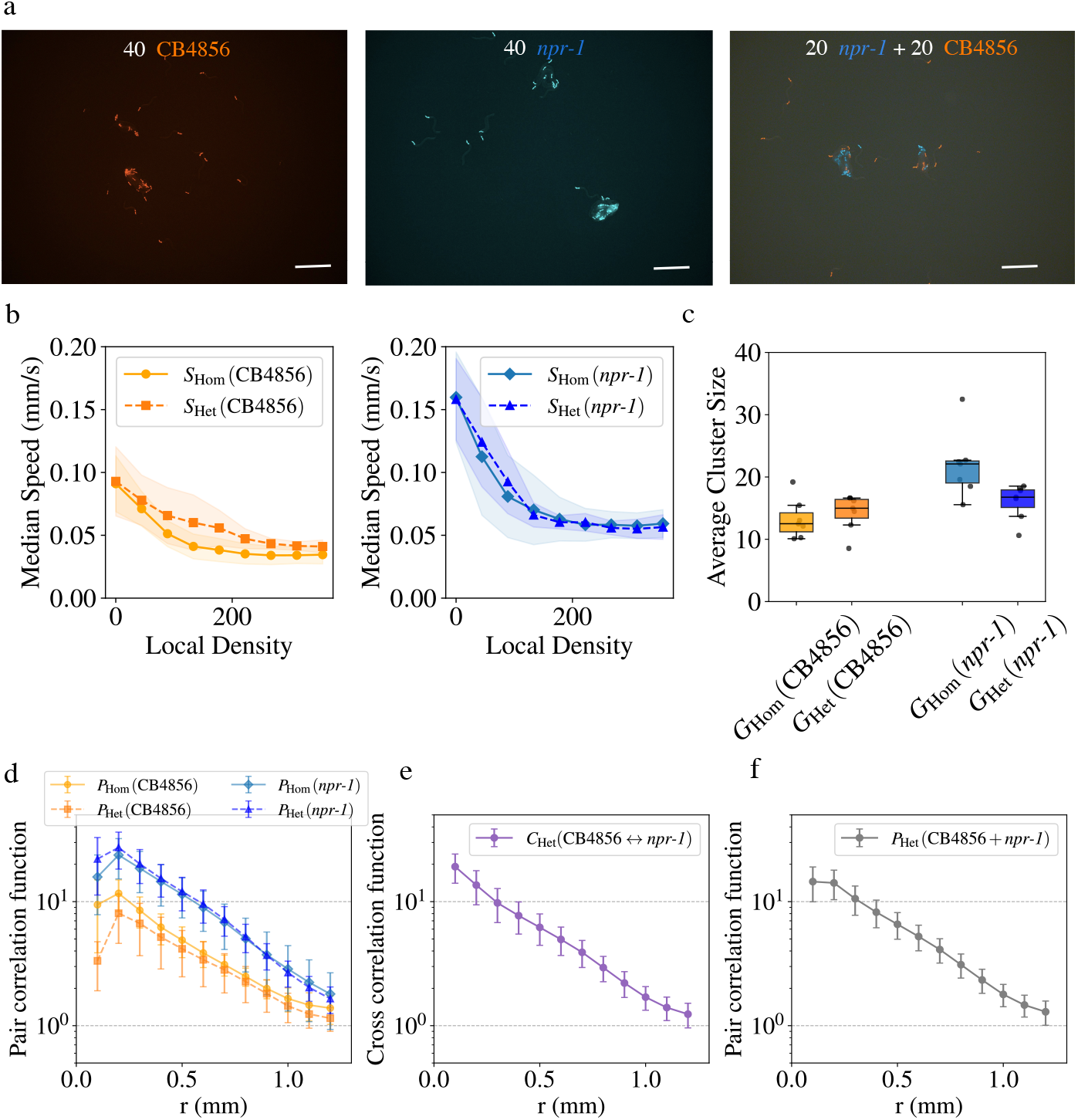
Behaviour quantification for MIX-2 experiments. **(a)** Representative snapshots of experiments of homogeneous experiments of 40 CB4856 individuals (left) or 40 *npr-1* individuals (middle), and heterogeneous experiment with 20 *npr-1* and 20 CB4856 individuals. Scale bars = 1 mm. See Supplementary Videos 2, 3, and 5 for additional representative examples of the behaviours. **(b)** Density dependent median speed (*S*) distributions for each strain in homogeneous (*S*_Hom_; solid line) and heterogeneous (*S*_Het_; dashed line) trials. The line shading represents the standard deviation of the data around the mean. **(c)** Average group size (*G*) for CB4856 and *npr-1* strains in homogeneous (*G*_Hom_) and heterogeneous (*G*_Het_) trials. The boxplots show the median (central line), interquartile range (box), and non-outlier data range (whiskers), with points beyond the whiskers representing outliers. **(d)** Pair correlation function (P) for the homogeneous groups (*P*_Hom_; solid line) in the heterogeneous groups (*P*_Het_; dashed line) for the CB4856 and *npr-1* strains. **(e)** Cross correlation function (*C*) from one strain type to the other. **(f)** Pair correlation function (*P*) considering both strains together in heterogeneous group. The error bars represent the the standard deviation of the data around the mean. The sample size is n = 7 for each experimental condition.

#### 2.2.1 Motility and density-dependence for each strain

Considering the genetic divergence between *npr-1* and CB4856, we next tested whether this affects motility response to local density in heterogeneous groups. Both strains exhibit decreased speed (*S*) with increasing local density, with the effect more pronounced in *npr-1* (Fig. 2b). These density-dependent trends are nearly identical in both homogeneous and heterogeneous groups, indicating that individual behavioural responses are robust to genetic heterogeneity in the social environment. Further analysis shows that these behavioural patterns hold regardless of whether the majority of local neighbours are of the same or the other strain; angular velocity (*W*) trends also show similar results (Supp. Fig. 2a). Thus, as in MIX-1, motility in MIX-2 is governed solely by local density, independent of the genetic identity of surrounding individuals. Since both strains in MIX-2 are aggregating, the decrease in speed with local density serves as a shared mechanism for aggregate formation in both strains.

#### 2.2.2 Cluster properties for each strain

In homogeneous trials, the average cluster size (*G*) reflects strain-specific aggregation patterns: both *npr-1* and CB4856 worms form aggregates, with *npr-1* forming larger clusters. In heterogeneous trials, the average cluster sizes fall between those observed in the respective homogeneous conditions (Fig. 2c): CB4856 worms are found in slightly larger clusters in the mixed condition than in their own homogeneous trials, suggesting that they join larger aggregates with *npr-1* worms, and conversely *npr-1* worms are found in the slightly smaller mixed aggregates. Looking at the fraction of individuals in aggregates (*F*), both *npr-1* and CB4856 show similar values across homogeneous and heterogeneous trials (Supp. Fig. 2b).

#### 2.2.3 Spatial organisation for each strain

Building on the cluster-level patterns, we examined the spatial organization of each strain in greater detail using the pair correlation function (*P*). As shown by the differences in *P*_Hom_, *npr-1* worms are consistently closer together than CB4856, in line with their larger cluster sizes, while CB4856 aggregates to a lesser extent (Fig. 2d). In heterogeneous groups, both *npr-1* and CB4856 retain their characteristic inter-individual spacing, as indicated by similar pair correlation values between homogeneous (*P*_Hom_) and heterogeneous (*P*_Het_) conditions (Fig. 2d). Mean neighbor distances also remain consistent across conditions for each strain (Supp. Fig. 2c). These results indicate that the presence of the other strain does not substantially alter strain-specific spacing or aggregation tendencies.

#### 2.2.4 Spatial relationship between strains

To understand how the two strains interact spatially in heterogeneous groups, we used the cross correlation *C*_Het_(CB4856 → *npr-1*). We found that *C* values were intermediate between the same-strain pair correlations (Fig. 2e), indicating spatial overlap. Comparing MIX-2 to MIX-1, the overall higher values of *C* reflect that CB4856 individuals in MIX-2 are more likely than N2 individuals in MIX-1 to be found within *npr-1* aggregates. This reflects a greater degree of spatial mixing in MIX-2, as further supported by the higher mean density overlap observed in MIX-2 compared to MIX-1 (Supp. Fig. 2d). The pair correlation function for all individuals in the heterogeneous group also has values that are intermediate relative to the homogeneous groups (Fig. 2f).

However, despite this increased overlap, the strains do not form uniformly spaced aggregates: each strain maintains its characteristic inter-individual spacing, as seen by similar pair correlation values in both homogeneous and heterogeneous conditions (Fig. 2d). The intermediate *C* values also suggest that *npr-1* worms are still more likely to be near other *npr-1* worms than CB4856, indicating that complete spatial mixing does not occur.

To summarize the MIX-2 trials with the distantly related strains *npr-1* and CB4856, we saw that while each strain maintains its characteristic aggregation behaviour, both strains readily form hybrid aggregates together. This demonstrates that behavioural compatibility, rather than genetic similarity, is sufficient for joint aggregation in *C. elegans*. These mixed aggregates arise from similar individual responses to local density but not from strain-specific interactions, highlighting that worms retain their intrinsic behavioural tendencies regardless of the genetic composition of their social environment.

### 2.3 Collective behaviour in heterogeneous *C. elegans* groups emerges from intrinsic strain-specific interaction mechanisms

We developed an agent-based simulation model to ask if strain-specific interaction rules can accurately reproduce the emergent spatial patterns from both homogeneous and mixed group experiments. While previous work has modelled worms as self-propelled filaments [11, 27], in our model we simplify this approach with a point-based representation to reduce complexity and improve flexibility for representing heterogeneous groups. Our model combines a modified random walk with social interactions (Fig 2a). The random walk component is designed to capture the motile behaviour of various *C. elegans* strains [28]. Social interactions are incorporated by a modification of the zonal model [5, 29]: to represent *C. elegans*, we introduce factors such that turning dynamics and speed are influenced by local neighbour positions and density (see Methods for details). To represent heterogeneous groups, we simulate a combination of two distinct types of agents, each of which is defined by a unique set of model parameters that reflect their individual behaviours in homogeneous social environments.

We chose values of basic model parameters based on previous work and in order to reflect the experimental configuration (Table 2), and then used a two-step procedure to set the values of key parameters that represent behavioural differences between strains. First, we fit model parameters for density-dependent speed using experimental trends for individual speed as a function of local density (Supp. Fig. 3a). In the model, differences in social responsiveness are represented by the social turning parameter, *α*. To capture experimentally observed differences between strains, we fit the value of *α* for each strain by matching the pair correlation function (*P*) and mean neighbour distance (M) in the homogeneous experiments to simulation results (Supp. Fig. 3b). With the fit parameter values for each respective strain, the model successfully reproduces the collective behaviour and spatial organization observed experimentally for all three strains in homogeneous groups (Fig. 3b, Supp. Fig. 3c, Supp. Vid. 6-8).

**Table 2:** Model parameters used in the simulations, along with their descriptions, units and values.

| Parameter | Description | Units | Value |
| --- | --- | --- | --- |
| $L$ | Arena size length | Distance ($\mu m$) | 10000 |
| $N$ | Total number of individuals | Non-dimensional | 40 |
| $t$ | Simulation length | Time steps | 500000 |
| $N_1$ | Number of type-1 individuals | Non-dimensional | Sim. dependent |
| $N_2$ | Number of type-2 individuals | Non-dimensional | Sim. dependent |
| $\mu_0$ | Speed average value at zero density | Distance/Time | Free parameter |
| $\mu_d$ | Speed density decay | Non-dimensional | Free parameter |
| $\mu_c$ | Speed baseline | Non-dimensional | Free parameter |
| $\tau_s$ | Speed auto-correlation time | Time (s) | 1.5 |
| $D_s$ | Speed diffusion constant | Distance<sup>2</sup>/Time ($\mu m^2/s$) | 200 |
| $\tau_w$ | Drift auto-correlation time | Time (s) | 0.1 |
| $D_w$ | Angular diffusion constant | Radians<sup>2</sup>/Time ($rad^2/s$) | 0.05 |
| $\tau_{fwd}$ | Forward run distribution constant | Time (s) | 50 |
| $\tau_{rev}$ | Reverse run distribution constant | Time (s) | 5 |
| $\alpha_i$ | Social turning responsiveness | Non-dimensional | Free parameter |
| $r_r$ | Repulsion zone radius | Distance ($\mu m$) | 100 |
| $r_a$ | Attraction zone radius | Distance ($\mu m$) | 800 |

**Table 3:** Model free parameters best-fit values that reproduce the behaviour of each simulated strain (denoted with an asterisk) in homogeneous groups.

| Parameter | N2* | CB4856* | *npr-1* * |
| --- | --- | --- | --- |
| $\alpha_i$ (Non dim.) | 0.09 | 0.13 | 0.30 |
| $\mu_0$ ($\mu m/s$) | 3.6 | 60.7 | 92.3 |
| $\mu_d$ (Non dim.) | 0.011 | 0.012 | 0.014 |
| $\mu_c$ ($\mu m/s$) | 42.6 | 31.7 | 56.1 |

**Figure 3:**
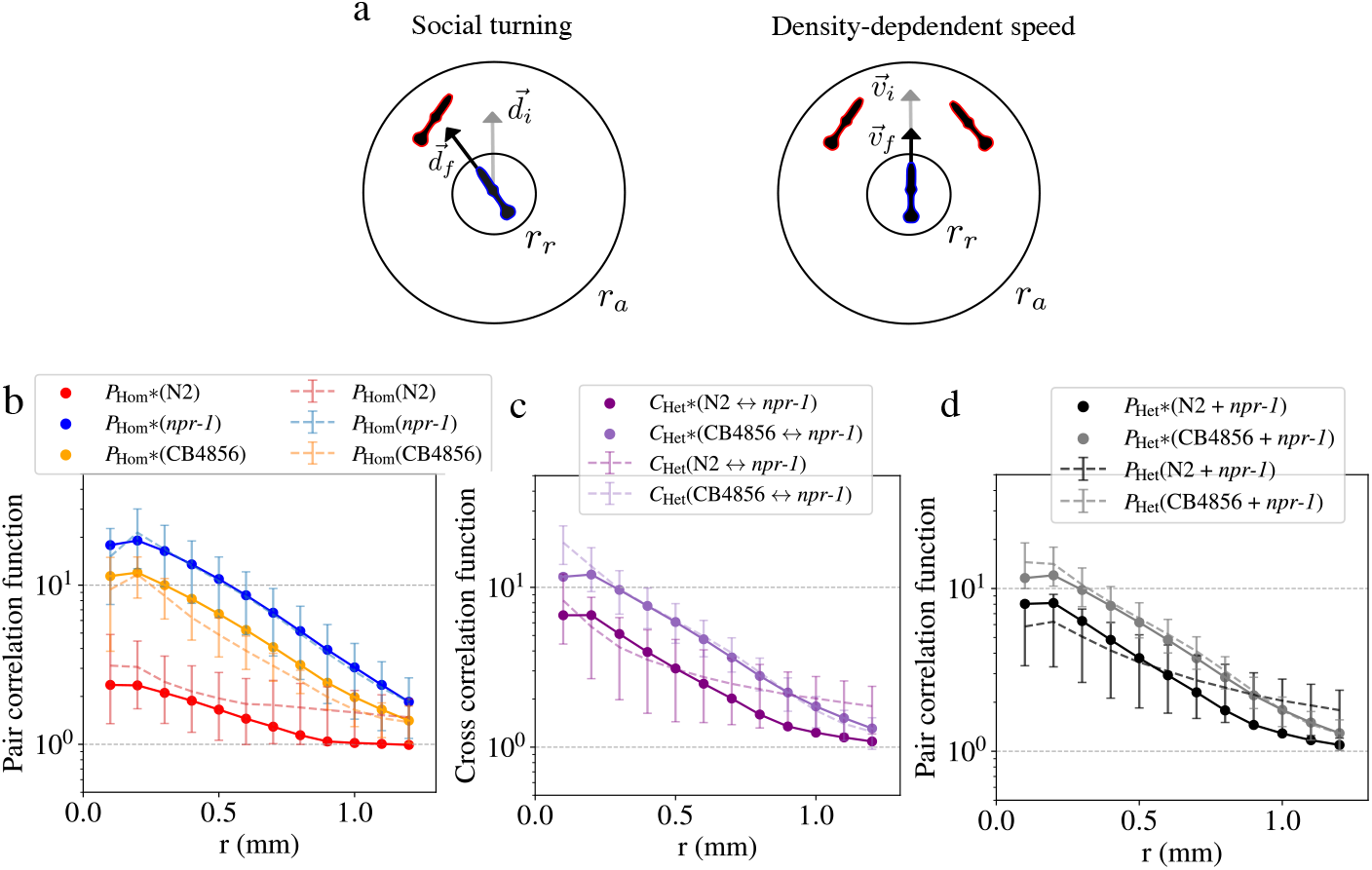
Individual-based simulation model. **(a)** Illustration of the social interactions in the model: Social turning and density-dependent speed. The inner circle represents the repulsion zone with radius *r*_*r*_ and the outer circle the attraction zone with radius *r*_*a*_ (Figure not drawn to scale). In the social turning schematic (left), 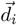 indicates the initial and 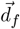 the final movement direction. In the density-dependent speed schematic (right), 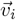 indicates the initial and 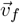 the final velocity. **(b)** Pair correlation function (*P*_*Hom*_) for homogeneous groups, comparing experiment (dashed line) and simulation model (solid line) results for each strain. Simulation results are indicated with an asterisk (*). **(c)** Cross correlation function (*C*) for the MIX cases, comparing experiment (dashed line) and simulations (solid line) results **(d)** Pair correlation function (*P*) for all individuals in the heterogeneous cases together, comparing experiment (dashed line) and simulations (solid line) results. The error bars represent the the standard deviation of the data around the mean.

To test the predictive ability of our model – which assumes indiscriminate interactions between strains in heterogeneous groups – we applied parameters derived from homogeneous trials directly to simulations of mixed groups. Both cross pair correlation (*C*) and mean density overlap (*D*) from the model closely match the experimental data (Fig. 3c, Supp. Fig. 3d,, Supp. Vid. 9-10). Additionally, the pair-correlation trends for overall spatial organization in heterogeneous groups are also well reproduced by the model (Fig. 3d).

These results demonstrate that the collective behaviour of heterogeneous groups of *C. elegans* can be predicted directly from intrinsic, strain-specific behaviours: the model, which uses parameters fit to experimental trends for homogeneous groups, accurately reproduces experimental outcomes for heterogeneous groups. This highlights how heterogenenous collective phenomena in *C. elegans* can emerge from simple, intrinsic behavioural rules, even in the absence of explicit recognition or selective interactions.

## 3. Discussion

This study examined the mechanistic foundations of how individual motion characteristics and inter-individual interactions influence the collective behaviour of explicitly defined heterogeneous groups of *C. elegans* on a shared food patch. We show that neither behavioural differences nor distant relatedness between members of the population produces detectable behavioural modulation of the individuals or the collective in heterogeneous social environments. In both MIX-1 and MIX-2 experiments, individual movement patterns and interaction mechanisms remain unchanged across homogeneous and heterogeneous conditions. Using a simulation model, we were able to accurately predict the behaviour of the heterogeneous groups directly from homogeneous group behaviour parameters inferred for each constituent strain. As individual worms in our heterogeneous group experiments behave only according to their own intrinsic motion and interaction characteristics, we conclude that composite collective phenotypes can arise in heterogeneous groups without requiring any individual-level behavioural modulation.

Multiple strains of *C. elegans* can co-occur in nature and share the same food patch [16, 17]. The apparent lack of behavioural modulation when behaviourally or genetically distinct strains occupy the same patch could in theory promote the co-existence of strains and the maintenance of diversity in this species by tolerating differences and, in light of MIX-1 results, even potentially reducing competition via differential spatial occupancy [30]. Despite the seemingly agnostic coexistence of different *C. elegans* strains under food abundance conditions, the situation might be different when food is depleted. We recently reported a novel collective dispersal behaviour called towering in multiple *Caenorhabditis* species including *C. elegans*, where many individuals physically writhe their bodies together to form a large tower structure to disperse together via hitchhiking [31]. In MIX-2 experiments we show that distantly related individuals readily form a physical hybrid aggregate together without discrimination. If this is also true under food depletion conditions, then all members of the *C. elegans* species in the vicinity should be able to build a larger tower together regardless of their genetic relatedness, which could potentially lead to better dispersal success and enhanced post-dispersal genetic diversity in newly colonised habitats. Whether heterogeneous groups of unrelated *C. elegans* individuals would physically congregate to tower together or start competing under resource depletion conditions is still an open question. Our work here provides the comparative context under resource rich conditions, as well as the conceptual and methodological framework to extend this comparison in the future to better understand the interaction mechanisms of heterogeneous collective behaviour in different resource and social environments.

Our finding of indiscriminate hybrid aggregate formation in *C. elegans* from MIX-2 experiments contrasts with the situation in another nematode species, *Pristionchus pacificus*, where kin-recognition plays a critical role in regulating social behaviour. Recent work has shown that while closely related strains of *P. pacificus* can form hybrid aggregates, distantly related strains form exclusive strain-specific clusters when placed on the same food patch [25]. This pattern is mediated by the biting mouth form and aggressive behaviours toward non-kin in this cannibalistic species, and is linked to the *self-1* kin recognition system [32]. Extending our work in the future to capture mechanistic details of selective interactions in these heterogeneous *P. pacificus* groups could provide a broader perspective on how recognition cues influence social interactions and group collective behaviour. By contrast, *C. elegans* lacks a known kin recognition system, and our findings suggest that aggregation in this species is driven by individual behavioural rules and not by strain-identity recognition. This result was surprising to us given the apparent genomic divergence [26] and expected pheromone profile differences [33] between the strains. Ascaroside pheromones are important signals that mediate communication and social interactions in nematodes [34, 35], but were shown not to be a regulator of aggregation behaviour in homogeneous *C. elegans* populations [11]. Here we find that between-strain pheromone differences in *C. elegans* are insufficient for identity discrimination in order to produce detectable behavioural modulation within the spatio-temporal context of our MIX-2 experiments.

Our work can be extended beyond *C. elegans* to a broader range of collective systems, providing a powerful methodology framework for investigating heterogeneity in various biological and artificial collective system. For example, we can extend our model to incorporate a selective interaction parameter to describe *P. pacificus* aggregation [25] or zebrafish shoaling [36] between distantly related individuals. While our study focused on behavioural differences, future research could explore additional dimensions of heterogeneity, such as differences in information access or physiological conditions. The methods we developed here can be applied to diverse systems, such as eusocial insect colonies with a division of labour between foragers and guards during foraging, fish schools with informed and uninformed members in decision-making, or even heterogeneous robotic swarms designed with different agent capabilities during task allocation. These further applications and extensions would deepen our understanding of how different forms of heterogeneity shape collective behaviour across biological and synthetic systems.

## 4. Materials and methods

### 4.1 *C. elegans* maintenance and strains

All *C. elegans* strains used in this work were maintained on standard nematode growth media (NGM) plates using standard protocol [37], and on a diet of *Escherichia coli* OP50. *C. elegans* worm culture and preparations were conducted under standard laboratory conditions at 20 °C. All strains used in this study can be found in the following table.

The SSD04 strain was created by introgressing the *aceIs1* red fluorescence marker into the CB4856 wild strain. This was achieved by crossing males of the ATU4301 strain with CB4856 hermaphrodites and selecting male progenies carrying the fluorescent marker for further backcrossing into the CB4856 background. The backcrossing and selection process was repeated ten times, and the marker was homozygouzed in the F10 generation.

### 4.2 Behavioural assays

All animals were cultured on NGM plates seeded with *E. coli* OP50 bacteria. Synchronized L1-diapause animals were obtained using a standard bleaching protocol [38], refed on seeded standard NGM plates with OP50 bacteria and incubated at 20°C for 65 ± 2 hours until they become Day-1 adults. To create a uniform food patch, a fresh overnight liquid culture *E. coli* OP50 was diluted in LB broth to obtain a OD_600_ = 0.6 *±* 0.2. Thirty minutes prior to starting the imaging of the experimental replicate, 3.5 cm diameter no-peptone NGM agar plates (standard NGM but with peptone removed to reduce bacterial growth during the experiment) were manually seeded with 20 *μ*L of OP50 bacteria. The droplet was left to dry at room temperature creating a 10 *±* 1 mm diameter circle of OP50 food arena. Fresh seeding helps to prevent the ring effect where there is a higher concentration of bacteria along the border of the food patch, as this is known to cause bordering behaviour in *C. elegans* that confounds the aggregation phenotype. Day-1 adults were washed off the culture NGM plates with 1 mL M9 buffer, washed twice more in M9 by centrifugation at 1500 rpm, and dispensed as small droplets onto the seeded imaging plate around the circumference of the food patch. A total of 40±4 worms for the homogeneous case and 20±2 for each type of worm in the heterogeneous case were dispensed. Note that for the heterogeneous case, the worms were not premixed; instead, the two strains were dispensed separately and randomly around the food patch. After the worms were dispensed, the droplets were allowed to evaporate (5-10 min) and the plate was gently vortexed for 10 seconds to randomly distribute the worms. Once more than 30 worms reached the food patch, the imaging plate was transferred to the behavioural microscope (details below) and the imaging commenced.

### 4.3 Data acquisition

In MIX-1 experiment worms were recorded for 1h and in MIX-2 experiments worms were recorded for 45 minutes, both at 10 fps. The first 10 minutes of each experiment were discarded during the analysis to account for the acclimation period, and the final 15 minutes of the MIX-1 experiments were discarded to match the experimental duration of MIX-2 for comparision between the two sets of experimental results. Imaging conditions were maintained at 19 ± 1°C. Imaging was performed using a ZEISS Axio Zoom.V16 microscope with the PlanApo Z 0.5x objective with a magnification of 20x, and raw imaging data was acquired with the ZEN 3.5 Pro software. A colour camera (Axiocam 712 color) was used to record the worms, enabling subsequent tracking and identification of strains based on fluorescence marker colour differences. After the data acquisition videos were exported in AVI format. Tracking was performed using the TRex software [22]; in the tracking software, the two colour channels (red and blue) were separated in a heterogeneous group to obtain the motion trajectories and strain identities of each strain. A total of 7 experimental replicates were obtained for each of the experiment condition.

### 4.4 Individual behaviour quantification

We quantified individual behaviour and responses to local density by analysing median speed (*S*) and angular velocity (*W*). Local density was defined as the number of individuals within one body length (1 mm) of the focal individual. To investigate the effects of group composition on behaviour, we applied a threshold in heterogeneous groups. Specifically, we examined local densities comprising 50% or more individuals of the same or the other type.

### 4.5 Collective behaviour quantification

To quantify the collective behaviour of the groups, we used position-based metrics and introduce measures that capture variations in the collective behaviour of homogeneous and heterogeneous groups across different sets of experiments, as well as the spatial relationships between different strain pairs in heterogeneous groups.

#### 4.5.1 Pair correlation function (*P*)

The pair correlation function, also known as the radial distribution function (typically described with the notation *g*(*r*)), is extensively utilized to characterize physical systems, and has been previously successful in capturing the differences in the collective behaviour between different *C. elegans* strains [11]. First we looked at the pair correlation function between all the pairs.

Aggregation was quantified using the pair correlation function, *P*, which quantifies the probability of finding another individual at a certain distance *r*. *P* is normalized by the number of individuals so that *P* = 1 represents no spatial organization structure (i.e., random distribution of particles), *P* > 1 represents aggregation, and *P* < 1 represents a lack of neighbours at a certain distance *r*.

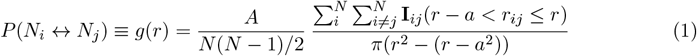

Where *A* is the arena size.

#### 4.5.2 Cross correlation function (*C*)

Then we introduced a modified version of the pair correlation function: the cross correlation function (*C*), used to characterize the relative positions of the two types of individuals, computing the distances between two different types of individuals (and vice-versa), named type-1 and type-2 for notation convenience.

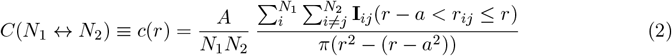

Where *N*_1_ and *N*_2_ are the total number of type-1 and type-2 individuals respectively, and *A* is the arena size.

#### 4.5.3 Mean neighbour distance (*M*)

We calculated the mean neighbour distance (*M*) by randomly sampling one agent from each frame and computing the distances to all other agents, from which we derive the mean.

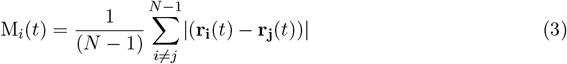

Subsequently, we averaged these mean distances across all frames in the experiment.

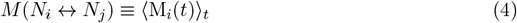

To compare the simulation and experimental results we introduce a scaling factor.

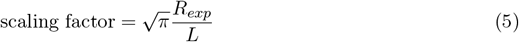

The simulation is run in a square arena with length size (*L*) with periodic boundary conditions, while experimental data comes from a circular arena of radius *R*_*exp*_. To compare M values from simulations and experiments, we scaled the simulated M by the ratio of the square root of the experimental area to the side length of the simulation arena. This ensures consistent density and accounts for differences in arena geometry.

#### 4.5.4 Mean density overlap (*D*)

To quantify the degree of overlap between the two types of individuals, we computed the mean coarse-grained density overlap (*D*) across the experiment. This metric is adapted from condensed matter physics where it has been used to quantify spatial segregation in densely packed (space-filling) systems, such as binary mixtures of particles or cell sheets [39]. Since worms do not form space-filling sheets, we adapted the metric to use density functions instead of Voronoi tesselation to better describe spatial overlap. First, we obtained the position distributions of both types of individuals and smooth them using a Gaussian filter. *A* Gaussian filter is a common smoothing technique in image and signal processing, where the parameter sigma (*σ*) controls the standard deviation of the Gaussian function. Larger sigma values result in stronger smoothing by distributing the weights over a broader region. We used *σ* = 1.5. Next, we computed the Hellinger distance (*D*_*H*_) between the two smoothed (or coarse-grained) position distributions at each time frame, denoted as *R*(*t*) and *Q*(*t*). The Hellinger distance quantifies the similarity between two probability distributions, with lower values indicating greater overlap and higher values signifying greater segregation. We computed this distance across all experimental frames, allowing us to track segregation dynamics over time.

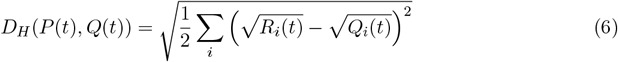

After computing the Hellinger distance for each time frame, we took the mean across all time frames to obtained an overall measure of segregation.

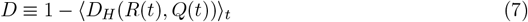

If the two types of individuals are completely segregated then D has a value of 0, whereas complete overlap is indicated by a value of 1. To establish a baseline for comparison we computed the mean density overlap randomizing the positions of the individuals.

#### 4.5.5 Fraction of individuals in clusters (*F*)

The previous metrics introduce a coarse-grained analysis, which do not provide information about the internal structure, composition, or number of clusters. Also, simple and coarse-grained cluster statistics have the drawback that cluster fission and fusion processes can alter the statistics, even when the individuals remain equally segregated. To address this, we performed a micro-scale level quantification. We used topological analysis to determine a representative value for the distance threshold used to define a cluster. This means the threshold is not chosen arbitrarily but instead emerges from the data. We computed the zeroth Betti number (*b*_0_), which is a topological invariant that represents the number of connected components in a topological space [40].

Essentially, we built the network by expanding the parameter *ϵ* (connection threshold) and identify a suitable interaction range by finding the intersection point of the high and low slope curves of the normalized zeroth Betti number (< *b*_0_/*N* >). This allowed us to estimate a characteristic length—approximately 0.4 mm (nearly half the body length of a individual). This value is based on data from the homogeneous *npr-1* experiments, which show the highest degree of aggregation (Fig. 4).

**Figure 4:**
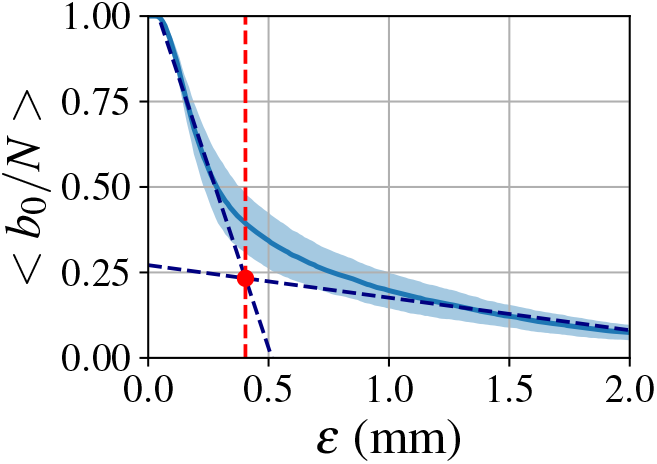
Setting the distance threshold for cluster analysis. Normalized zeroth Betti number (< *b*_0_/*N* >) in function of the connection threshold *ϵ* (mm). The red dot indicates the intersection point (*ϵ* = 0.4 mm) between the fitted high and low slope curves.

We then defined a cluster as a connected component in a network considering all the individuals with a connection threshold of *ϵ* = 0.4 mm and a minimum of three individuals. Once clusters were defined we could compute the fraction of individuals of each strain that are inside one of them. Then we computed the mean of this value across each replicate.

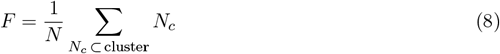

Where *N* is the total number of individuals and *N*_*c*_ is the number of individuals that belong to a cluster. T

#### 4.5.6 Average cluster size (*G*)

Using the same cluster definition as in the previous section allows us to quantify the (weighted) average cluster size (*G*) that an individual worm belongs to; In other words it gives the probability of finding an individual in a cluster of certain size. Using a distance threshold, we defined a network in which worms are considered “connected” if they lie within this threshold. A cluster was then defined as a connected component within this network that includes a minimum of three individuals. This approach allows us to quantify the average cluster size (*G*) that an individual worm belongs to. For example in a cluster of 40 individuals, lower values represents more solitary behaviour, while G = 40 would represent all worms in a trial always remaining in a single aggregated cluster.

Weighting by cluster size gives greater importance to individuals in larger clusters, making the metric more representative of collective aggregation patterns when cluster sizes are unevenly distributed. As a result, isolated individuals contribute less to the overall measure.

After visual inspection, we found that strong overlap between worms within aggregates caused some individuals to be missed during tracking. To correct for this in the metric—which is not normalized by the total number of individuals— we accounted for the known group size of 40 worms in the arena by adding a correction step that randomly distributes the missing individuals across existing clusters.

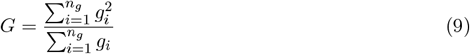

Where *N*_*g*_ is the number of components and *G*_*i*_ is the number of individuals in that component. Then we computed the mean of this value across each replicate.

### 4.6 Individual-based model

We implemented an individual-based model in order to gain further insights into how and which basic behavioural rules influence the worm collective behaviour in both homogeneous and heterogeneous groups. The model incorporates two different types of agents interacting in a square arena with size length L and periodic boundary conditions. The model combines elements from previous work studying *C. elegans* and fish behaviour [28, 29]. Furthermore, it considers independent dynamics in the speeds, turning and reversal events.

The translation motion of the individuals can be described as a change of the centroid velocity **v**(**t**) which can be decomposes into speed *s*(*t*) and direction of motion *ϕ*(t):

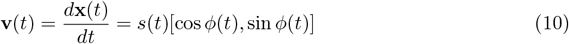

The direction of motion of the simulated worms can be described as.

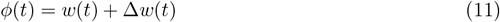

We defined the orientation of individuals *w*(*t*) by the centroid to head direction capturing the turning dynamics and Δ*w*(*t*) describing the forward and reverse states of motion. We simulated a total of number of agents *N* = *N*_1_ +*N*_2_, where *N*_1_ and *N*_2_ are the number of type-1 and type-2 individuals, respectively. Each type possesses a distinct set of model parameters, enabling us to simulate various agent types based on their motility and interaction rules.

### 4.6.1 Diffusive turning with drift

We implemented a simplified version of the zonal model where we considered repulsion and attraction zones. The two social zones, repulsion and attraction, are defined by the distance radius *r*_*r*_ and *r*_*a*_ respectively. For a focal individual *i* and one of its neighbours *j*, the distance between the two is *r*_*ij*_ = |**r**_**j**_ − **r**_**i**_|. The preferred motion direction from the zonal model is determined as:

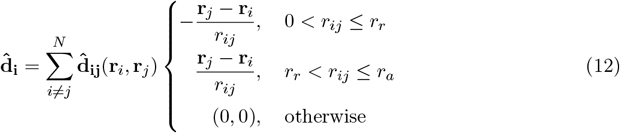

This direction is used to calculate an effective social torque:

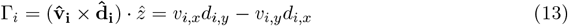

Where 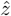 is the unit vector in the z-direction. The orientation dynamics are captured by a simple model that combines the previously introduced social interactions, with drift and stochastic diffusion:

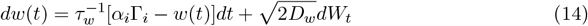

Where *α*_*i*_ is the social coupling strength or ‘turning responsiveness’, *τ*_*w*_ the relaxation time, and the random fluctuations arise from a Wiener noise process, dW_*t*_, with magnitude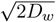.

### 4.6.2 Density dependent speed dynamics

The speed dynamics of each individual are described by an Ornstein-Uhlenbeck stochastic process:

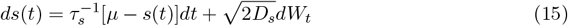

Which describes random fluctuations arising from a Wiener noise process, dW_*t*_, with magnitude 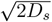, and relaxes with a time scale *τ*_*s*_ to an average value of *μ*_*s*_ = ⟨s ⟩.

Considering previous empirical observations, we incorporated a density-dependent speed, where the average speed decreases exponentially with the local density, plus an additional term.

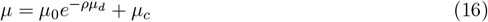

We defined the speed at zero density as *μ*_0_ (individuals without any neighbour inside their attraction zone), and *μ*_*c*_ as the baseline speed. Speed decays exponentially with an exponent *μ*_*d*_, and *ρ* indicates the local density of worms inside the attraction zone of radius *r*_*a*_ surrounding an individual.

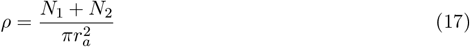

Where *N*_1_ and *N*_2_ are the number of type-1 and type-2 individuals respectively.

### 4.6.3 Forward and reverse turns

We described the forward and reverse runs, implying a sudden change in the direction Δ*w*(*t*) = 180^*°*^, (or a ‘random telegraph process’), a stochastic process characterized by sudden, random switches between two distinct states:

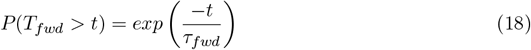

and,

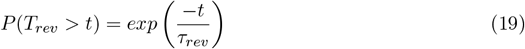

The time distributions of forward and reverse turns intervals are (*T*_*fwd*_ and *T*_*rev*_), and are determined by the coefficients (*τ*_*fwd*_ and *τ*_*rev*_).

#### 4.6.4 Model parameters

Fixed motility parameters are set to represent characteristic *C. elegans* movement patterns. Since the model is an abstraction, parameters are set according to the behavioural characteristics and the temporal and spatial scales of *C. elegans* behaviour. The distance parameters (attraction and repulsion zone radius) are determined based on the peak and decay of the pair correlation function, while also accounting for short-range interactions between individuals.

#### 4.6.5 Parameter fitting

Social responsive and speed-related parameters are selected to represent different strains and to highlight key differences. To determine the value of the social turning responsiveness parameter *α* for the model that better matches the data, we defined a distance function, DIST_exp-sim_, that quantifies the difference between the pair correlation functions obtained from experiments and homogeneous simulations. Specifically, we used the euclidean norm of the logarithmic differences between the experimental and simulated pair correlation function and mean neighbour distance.

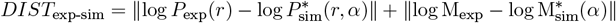

Where *P*_exp_(*r*) and 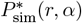 represent the experimental and simulated pair correlation functions, respectively, as functions of inter-individual distance *r* (analogous for the mean neighbour distance (M)). The parameter *α* is chosen to minimize DIST_exp-sim_, ensuring the best match between experimental and simulated data. Taking the logarithm of the pair correlation function helps to emphasize differences across different scales, particularly in cases where *P*(*r*) spans within two orders of magnitude. We incorporated an interpolation on the simulation results to have finer grid and better estimate the best fit parameters. Speed-related parameters are fitted with an exponential curve based on the median speed-density decay from the data (Supp. Fig. 3b).

## 5. Supplementary Material

Supplementary experimental and simulation results videos:

- Video 1 - Sample homogeneous N2 experiment
- Video 2 - Sample homogeneous *npr-1* experiment
- Video 3 - Sample homogeneous CB4856 experiment
- Video 4 - Sample heterogeneous MIX-1 (N2 + *npr-1*) experiment
- Video 5 - Sample heterogeneous MIX-2 (CB4856 + *npr-1*) experiment
- Video 6 - Sample homogeneous N2 simulation
- Video 7 - Sample homogeneous *npr-1* simulation
- Video 8 - Sample homogeneous CB4856 simulation
- Video 9 - Sample heterogeneous MIX-1 (N2 + *npr-1*) simulation
- Video 10 - Sample heterogeneous MIX-2 (CB4856 + *npr-1*) simulation

## 6. Data and code availability

Datasets used in this study will be available on Zenodo upon publication, which includes original tracked data from the experiments in this study. Codes for analyses, modelling and generating figures are available on GitHub in the following link: github.com/SerenaDingLab/Font-Massot_et_al_WORMIX

## 7. Acknowledgments and funding

We would like to credit Iris Bernstein for help with data collection and animal maintenance. We would like to thank Jayme Weglarski for laboratory technical support and media preparation. This study was funded through the Deutsche Forschungsgemeinschaft (DFG, German Research Foundation) under Germany’s Excellence Strategy EXC 2117 – 422037984, the Max Planck Institute of Animal Behavior, the International Max Planck Research School for Quantitative Behavior, Ecology & Evolution, and the BABOTs consortium grant (Horizon Europe, European Innovation Council Pathfinder Work Programme under grant agreement no. 101098722).

## 8. Author contributions

- N.F-M. : Conceptualization, Data curation, Formal analysis, Investigation, Methodology, Software, Visualization, Writing – original draft, Writing – review & editing
- J.D.D. : Conceptualization, Formal analysis, Investigation, Methodology, Software, Supervision, Validation, Visualization, Writing – original draft, Writing – review & editing
- S.S.D. : Conceptualization, Funding acquisition, Investigation, Methodology, Project administration, Resources, Supervision, Validation, Writing – original draft, Writing – review & editing

