## Supplementary Information for "Intrinsic strain-specific behaviour predicts emergent collective aggregation in heterogeneous *C. elegans* groups"

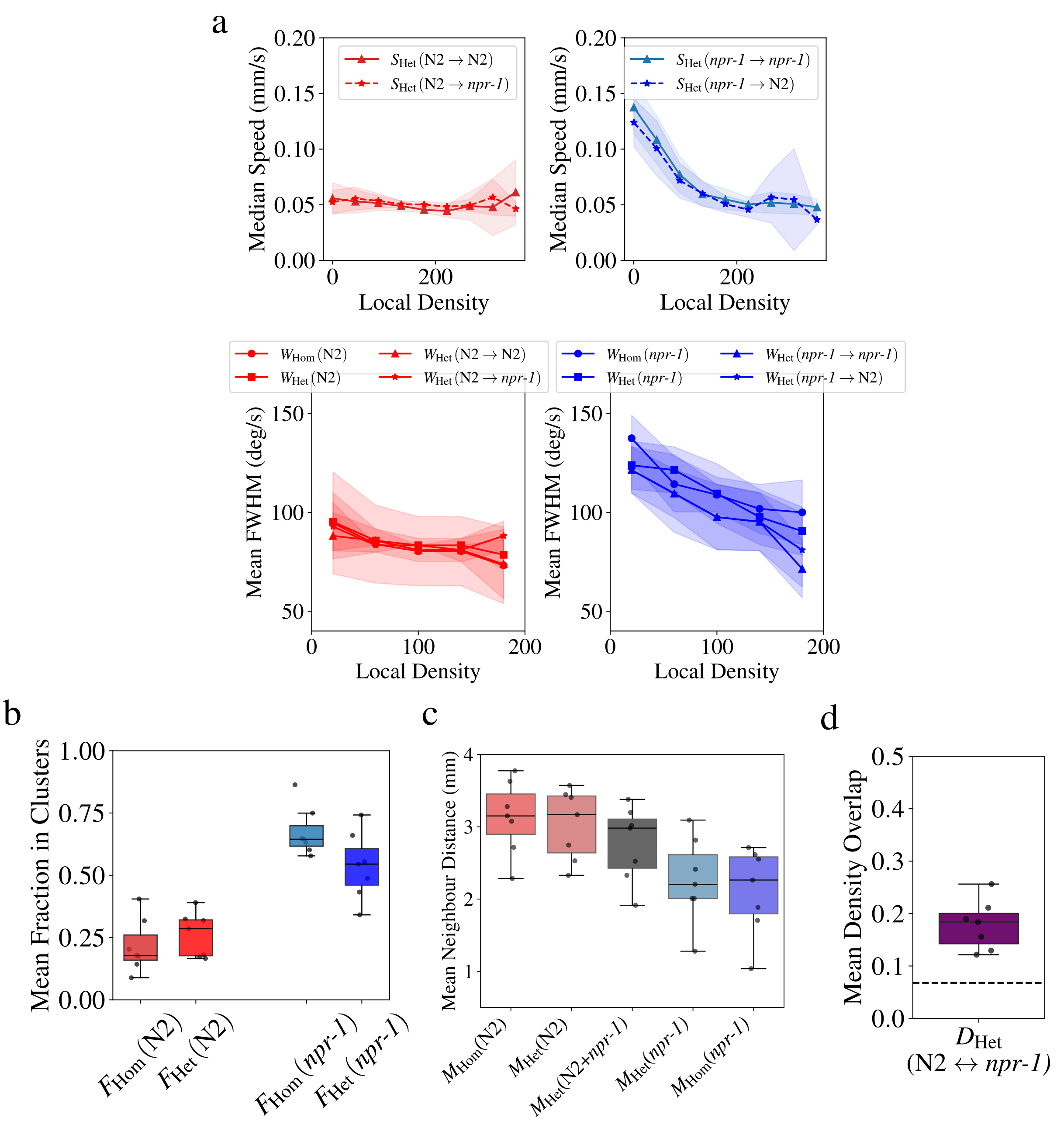


Supplementary Figure 1: **Additional metrics for MIX-1.** **(a)** Density dependent median speed ($S$) and mean FWHM (Full width at half maximum) of the angular velocity distributions ($W$) for the N2 and *npr-1* strains in homogeneous and heterogeneous environments. The arrows indicate the relation between the selected focal individual and neighbour type for each metric. **(b)** Mean fraction of individuals in clusters ($F$) for N2 and *npr-1* strains in homogeneous ($F_{\text{Hom}}$) and heterogeneous ($F_{\text{Het}}$) groups. **(c)** Mean neighbour distance ($M$) for the homogeneous groups, heterogeneous groups considering each strain separately, and considering both strains at the same time $M_{\text{Het}}(\text{N2} + \textit{npr-1})$ for N2 and *npr-1* **(d)** Mean density overlap ($D_{\text{Het}}$) between the N2 and *npr-1* strains. The sample size is $n = 7$ for each experimental condition.

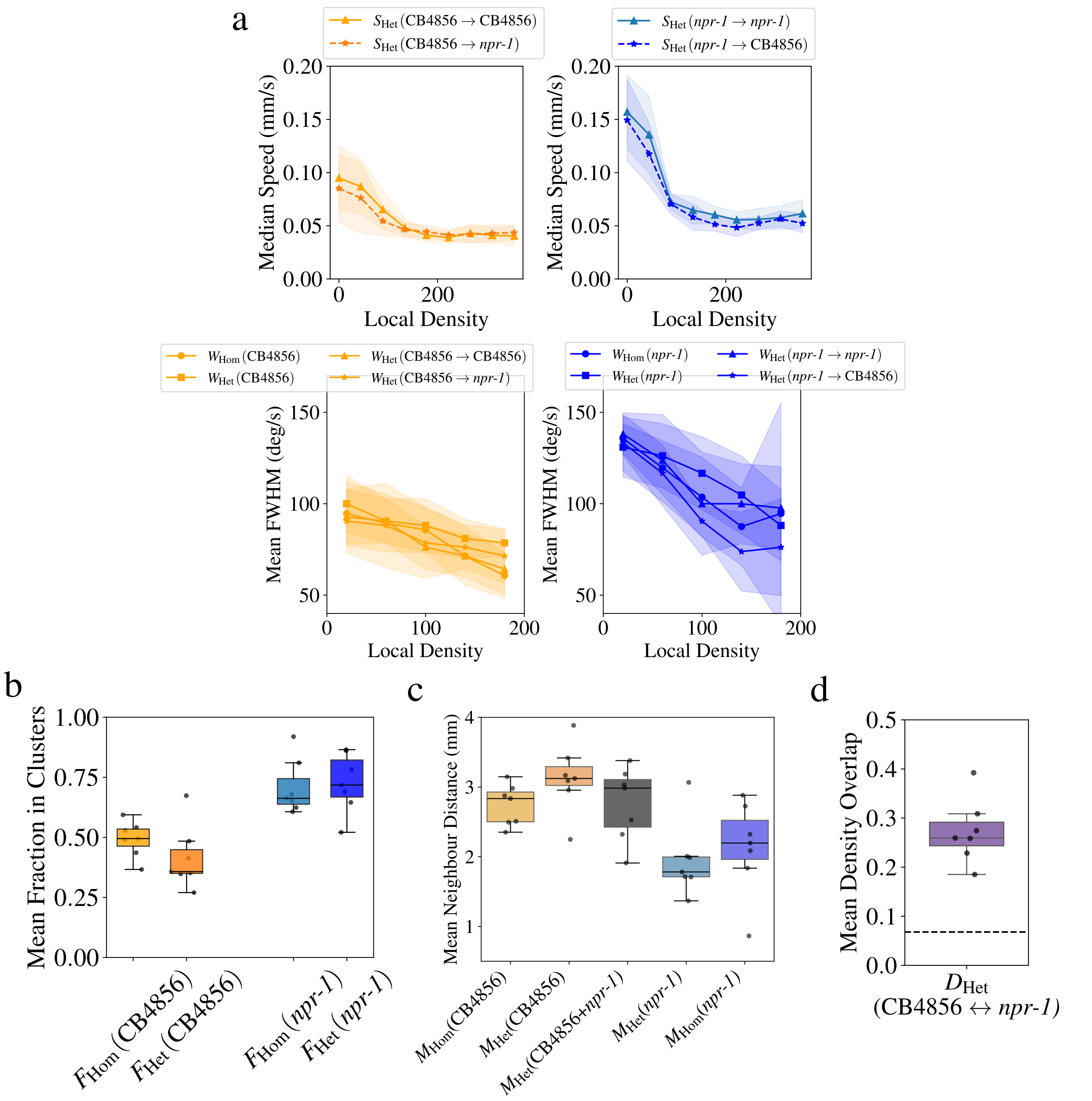


Supplementary Figure 2: **Additional metrics for MIX-2.** **(a)** Density dependent median speed ($S$) and mean FWHM (Full width at half maximum) of the angular velocity distributions ($W$) for the CB4856 and *npr-1* strains in homogeneous and heterogeneous environments. The arrow indicates the relation between the selected focal individual and neighbour type for each metric. **(b)** Mean fraction of individuals in clusters ($F$) for CB4856 and *npr-1* strains in homogeneous ($F_{\text{Hom}}$) and heterogeneous ($F_{\text{Het}}$) groups. **(c)** Mean neighbour distance ($M$) for the homogeneous groups, heterogeneous groups considering each strain seperatelly, and considering both strains at the same time $M_{\text{Het}}(\text{CB4856} + npr\text{-}1)$ for CB4856 and *npr-1* **(d)** Mean density overlap ($D_{\text{Het}}$) between the CB4856 and *npr-1* strains. The sample size is $n = 7$ for each experimental condition.

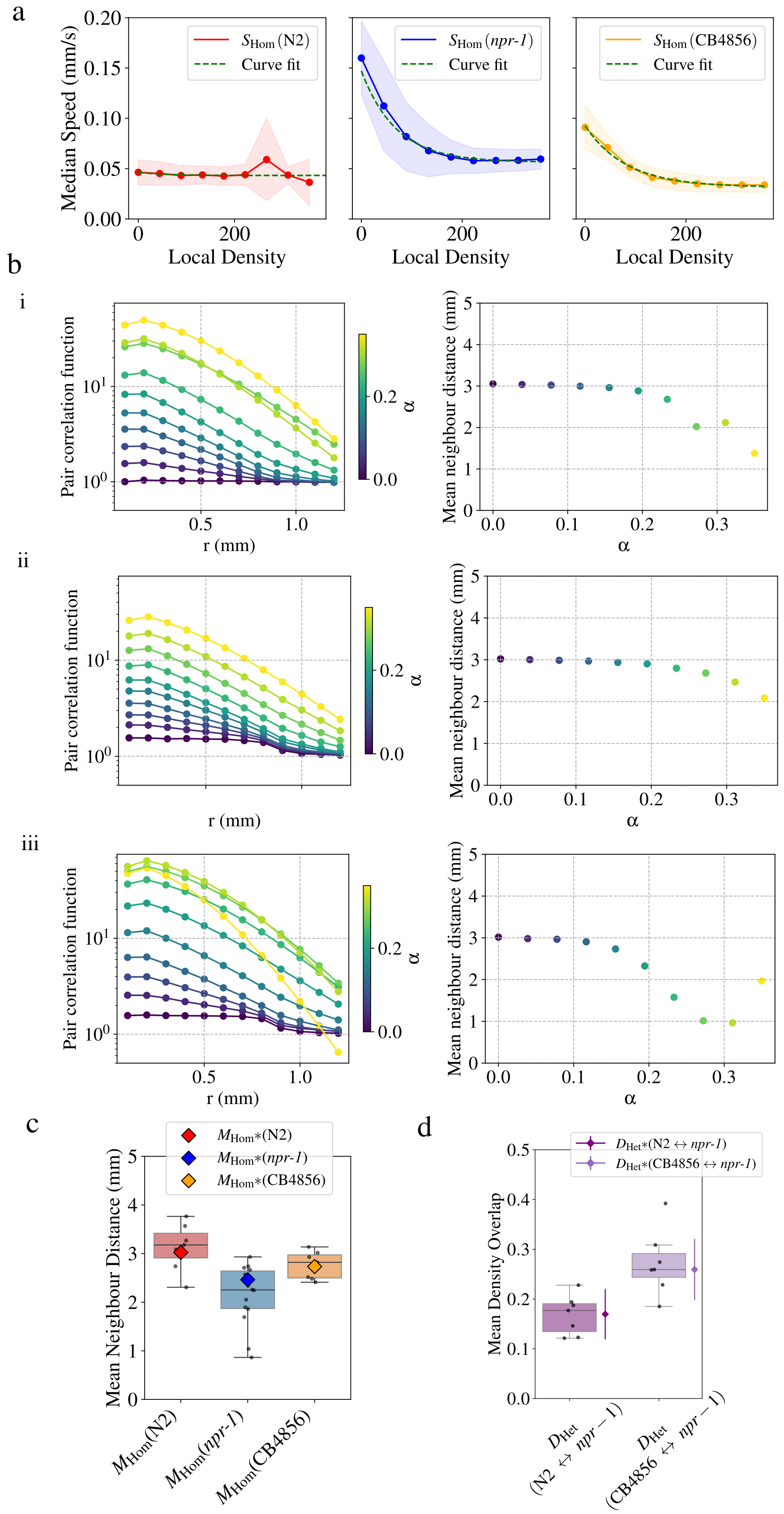


Supplementary Figure 3: **Simulation parameter fitting and additional simulation results.** **(a)** Fitted exponential curves for the density dependent median speed ($S$) vs the local density for the three studies strains (N2, *npr-1*, CB4856) **(b)** Exploration of the parameter $\alpha$, which sets the strength of social turning interactions. Shown are the pair correlation function ($P$) and mean neighbour distance ($M$), simulated using speed parameters set for the different strains: i) N2, ii) *npr-1* and iii) CB5856. **(c-d)** Comparison experimental and simulation results using best fit parameters (Table 3), showing **(c)** mean neighbour distance ($M$) and **(d)** mean density overlap ($D$). Error-bars are included to account for the fluctuations in the metric. Simulations results are indicated with an asterisk (*).
